# Neurophysiological and Autonomic Dynamics of Threat Processing During Sustained Social Fear Generalization

**DOI:** 10.1101/2024.04.16.589830

**Authors:** Jourdan J. Pouliot, Richard T. Ward, Caitlin M. Traiser, Payton Chiasson, Faith E. Gilbert, Andreas Keil

**Author notes:** Corresponding author: Jourdan J. Pouliot Department of Psychology Center for the Study of Emotion & Attention University of Florida PO Box 112766 Gainesville, FL 32611. denotes joint first authorship (Jourdan J. Pouliot, Richard T. Ward).

## Abstract

Survival in dynamic environments requires that organisms learn to predict danger from situational cues. One key facet of threat prediction is generalization from a predictive cue to similar cues, ensuring that a cue-outcome contingency is applied beyond the original learning environment. Generalization has been observed in laboratory studies of aversive conditioning: behavioral and physiological processes generalize responses from a stimulus paired with threat (the CS+) to unpaired stimuli, with response magnitudes varying with CS+ similarity. In contrast, work focusing on sensory responses in visual cortex has found a sharpening pattern, in which responses to stimuli closely resembling the CS+ are maximally suppressed, potentially reflecting lateral inhibitory interactions with the CS+ representation. Originally demonstrated with simple visual cues, changes in visuocortical tuning have also been observed in threat generalization learning across facial identities. It is unclear to what extent these visuocortical changes represent transient or sustained effects and if generalization learning requires prior conditioning to the CS+. The present study addressed these questions using EEG and pupillometry in an aversive generalization paradigm involving hundreds of trials using a gradient of facial identities. Visuocortical ssVEP sharpening occurred after dozens of trials of generalization learning without prior differential conditioning, but diminished as learning continued. By contrast, generalization of alpha power suppression, pupil dilation, and self-reported valence and arousal was seen throughout the experiment. Findings are consistent with threat processing models emphasizing the role of changing visucocortical and attentional dynamics when forming, curating, and shaping fear memories as observers continue learning about stimulus-outcome contingencies.

## Introduction

The ability to generalize learned responses to threat is a necessary component of adaptive cognition. In naturalistic conditions, the mammalian visual system is tasked with parsing complex, changing environments for possible sources of danger (Li & Keil, 2023). To do so adaptively, stimulus-outcome contingencies cannot be mapped one-to-one but must flexibly envelope both the encountered threat and stimuli that are close in resemblance (Averbeck & Costa, 2017; Lissek et al., 2014). This serves an important function, because it allows mammals to form predictions about novel stimuli or contexts when gathering resources or avoiding predators (Fanselow, 1994). Dysregulation of this process is considered an etiological factor to the development and maintenance of anxiety disorders, in which individuals are thought to overgeneralize threat responses to harmless stimuli (Lissek et al., 2014). Developing a precise model of fear generalization and threat discrimination has implications for clinical work as well as in the field of artificial intelligence, where optimal generalization procedures are central to classification algorithms (Mitchell et al., 1986; Ludvig & Koop, 2008).

A substantial body of work has used Pavlovian aversive conditioning paradigms to investigate the physiological processes accompanying aversive generalization learning. These paradigms induce fear generalization by pairing an aversive unconditioned stimulus (US; e.g. an electric shock or loud noise) with a neutral stimulus (e.g. shapes, gratings, faces), such that presentation of the conditioned stimulus (CS+) alone elicits an aversive response in absence of the US. Another stimulus (CS–) is never paired with the US. Typically, after this initial period of acquisition, a generalization phase follows in which generalization stimuli (GSs) are presented. GSs systematically vary in their similarity to the CS+ along a gradient of features such as size, orientation, color, or location (Antov et al. 2020; Friedl & Keil 2021). Although the GSs are never paired with a US, they tend to elicit heightened defensive responses (e.g., Friedl & Keil, 2020; Lissek et al., 2014). These responses typically diminish monotonically, with decreasing similarity to the CS+ (McTeague et al., 2015), resulting in a so-called generalization tuning pattern (see figure 1, left panel). Such patterns have been found for a wide range of responses, including self-reported affect as well as somato-visceral measures such as skin conductance, pupil dilation, and heart rate change (Ahrens et al., 2016). In the absence of any generalization, all-or-nothing patterns (figure 1, middle panel) may emerge, showing selective heightening for the CS+ only, with no enhancement for any of the GSs. This type of pattern has, for example, been observed in some rodent studies with compound conditioned cues (Mclaren & Mackintosh, 2002) and in studies of human auditory conditioning (Farkas et al., 2023; Heim & Keil, 2006).

**Figure 1.**
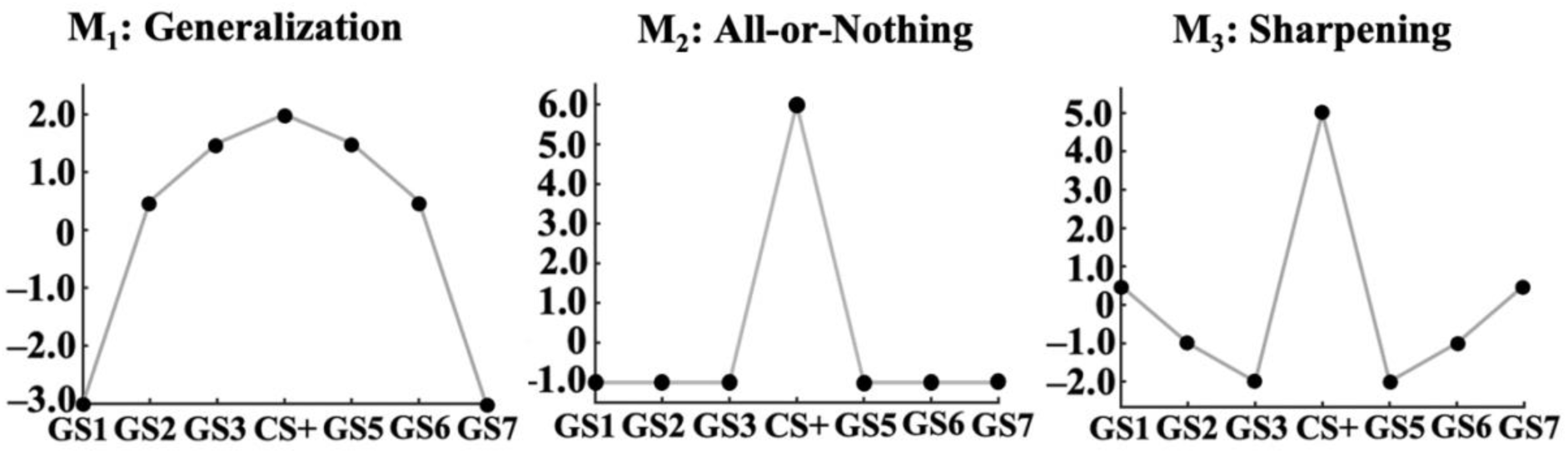
Hypothetical patterns emerging during generalization learning. Left: Quadratic trend showing a broad generalization pattern. Middle: discrimination of only the CS+. Right: Difference-of-Gaussians/sharpening pattern with suppression of responses to GSs with high similarity to the CS+.

At the level of population responses in the human brain, electrophysiological and hemodynamic imaging studies have converged to characterize a large-scale network of cortical and subcortical regions which likewise show generalization patterns (Dunsmoor & Paz, 2015). These regions include sensory cortices, in which neural populations display enhanced responses to CS+ stimuli (Skrandies & Jedynak, 2000; Heim & Keil, 2006). In human vision, several studies have used the steady-state visual evoked potential (ssVEP) to examine visuocortical activity during generalization learning at the population level. The ssVEP is an oscillatory response to stimuli that are regularly modulated in luminance or contrast at a fixed temporal rate (see Wieser et al., 2016 for review). It is readily quantified in the frequency domain, as a large oscillatory response with pronounced spectral amplitude at the same temporal frequency as the driving stimulus. These studies have converged to show that visuocortical areas encode threat-related information (Miskovic & Keil, 2013; Li & Keil, 2023), showing selective amplitude enhancement when viewing the CS+, compared to control stimuli such as CS–. Concurrent ssVEP-fMRI recordings suggest that CS+-related amplification of the ssVEP results from re-entrant feedback from the same anterior regions that have been characterized in a large body of hemodynamic imaging studies. In addition to higher-order visuocortical areas, these regions include the amygdala, the anterior cingulate cortex (ACC), and frontoparietal cortices often associated with attention control, such as the middle frontal gyrus (Petro et al., 2017; Uddin et al., 2019).

Visuocortical responses measured with ssVEPs often display a sharpening pattern (figure 1, right panel), showing preference for the CS+ accompanied by suppressed responses to the most similar GSs (McTeague et al., 2015; Antov et al., 2020). This may indicate inhibitory interactions between feature-specific sensory neurons that are located proximally to the neurons representing the CS+ feature (Li and Keil, 2023). Recent work by Stegmann and colleagues (2020) found sharpened tuning after initial differential conditioning with a gradient defined by facial identities, which was particularly pronounced in observers high in social anxiety. The present study builds on these findings, examining the time course of generalization learning across similar facial identities that is sustained of hundreds of trials, a laboratory equivalent of prolonged engagement with a given facial identity or person.

Stimulus-induced reduction of spectral power in the alpha band (8-13 Hz) has long been utilized as an index of an observer’s engagement with the visual world (Adrian & Matthews, 1934; Klimesch, 2012). Recently, transient alpha-power reduction has also been found to be sensitive to aversive conditioning (Panitz et al., 2019). For example, Bacigalupo and Luck (2022) found enhanced and longer parieto-occipital alpha suppression following presentation of the CS+ relative to the CS–.

### The present study

The present study expands on earlier work on aversive generalization learning with facial stimuli (Stegmann et al., 2020). Here, we examine the overarching hypothesis that visuocortical tuning functions measured with ssVEPs remain malleable as learning progresses beyond the initial conditioning and generalization test (Li & Keil, 2023). In addition to ssVEPs, we examined alpha-power changes and pupil dilation. Pupil dilation responses have been modulated by emotional arousal (Steinhauer et al., 2004; Bradley et al, 2008). During fear generalization, pupillary dilation follows a generalization pattern in response to a gradient of Gabor patches (Roesmann et al., 2022). The present work expected to replicate these findings with facial cues, allowing us to obtain a metric for establishing the extent to which observers showed evidence of sympathetic mobilization. It was hypothesized that visuocortical ssVEP patterns would follow a lateral inhibition gradient initially, potentially followed by heightened sharpening, generalization, or transition into an all-nothing-model. For parietal alpha power reduction, it was hypothesized that responses would follow a broad generalization gradient. These patterns were expected to be less pronounced after sustained acquisition (i.e. in later trials), reflecting transition to an all-nothing model as observed in auditory generalization condition, which likewise involves hundreds of trials (Farkas et al., 2023).

## Materials and Methods

### Participants

41 students (34 female) from the University of Florida participated in this study either for course credit or $70 monetary compensation. Participants were at least 18 years old and were screened for uncorrected visual impairments and epilepsy or family history of epilepsy. 3 participants withdrew from the study. Behavioral ratings from all participants who completed the study were used for analysis (N = 38 (31 female), M_AGE_ = 21.08, SD_AGE_ = 5.34). Of these, 31 participants identified as White (82.58%), 2 identified as Asian (5.26%), 1 identified as Black (2.63%), and 4 identified as Mixed Race/Other (10.53%). Of those who completed the study, 5 participants were excluded from EEG data analysis due to excessive noise in EEG data, leaving 33 participants (27 female, M_AGE_ = 21.12, SD_AGE_ = 5.61). Of those included in the EEG analyses, 26 participants identified as White (78.79%), 1 identified as Black (3.03%), 2 identified as Asian (6.06%), 4 identified as Mixed Race/Other (12.12%). Additionally, 4 participants who completed the study were excluded from pupil response analyses due to excessive noise, leaving 34 participants (29 female, M_AGE_ = 20.85, SD_AGE_ = 5.06). Of those included in pupil response analyses, 28 participants identified as White (82.35%), 1 identified as Black (2.94%), 1 identified as Asian (2.94%), 4 identified as Mixed Race/Other (11.76%). This study was approved by the local Institutional Review Board and was conducted in accordance with the Declaration of Helsinki.

Participants were part of a larger sample in a pre-registered study (https://osf.io/quy47) examining the effects of social anxiety disorder (SAD) on aversive generalization of facial stimuli, in which some participants were recruited for high scores on the Liebowitz Social Anxiety Scale (LSAS) prior to the task (Liebowitz et al., 1987). The sample utilized for the present study was drawn from the bottom half of the larger distribution of LSAS scores, excluding highly socially-anxious individuals, to minimize the effects of social anxiety symptoms.

### Materials and Stimuli

The present aversive generalization paradigm manipulated facial identity using morphing software to generate a feature gradient. The gradient stimuli (GSs) included seven monochrome female faces shown against a dark gray background. The initial facial stimuli were obtained from the Karolinska Directed Emotional Faces (KDEF) database (figure 2). Four of the seven faces were morphs of the middle face (the CS+) and one of the other original faces (GS1 or GS7), and were generated using open *MorphX* software (http://www.norrkross.com/). To correct for differences in contrast, a Gaussian filter with a full-width half maximum (FWHM) of 50 grayscale units was applied to all facial stimuli. Visual stimuli were shown on a calibrated LCD monitor (Display ++, Cambridge Research systems) with a refresh rate of 120Hz. The US was a burst of 92 dB(A) white noise presented from two speakers placed bilaterally behind the participant. The experiment was programmed and controlled using the MATLAB Psychophysics Toolbox (Brainard, 1997; Pelli, 1997).

**Figure 2.**
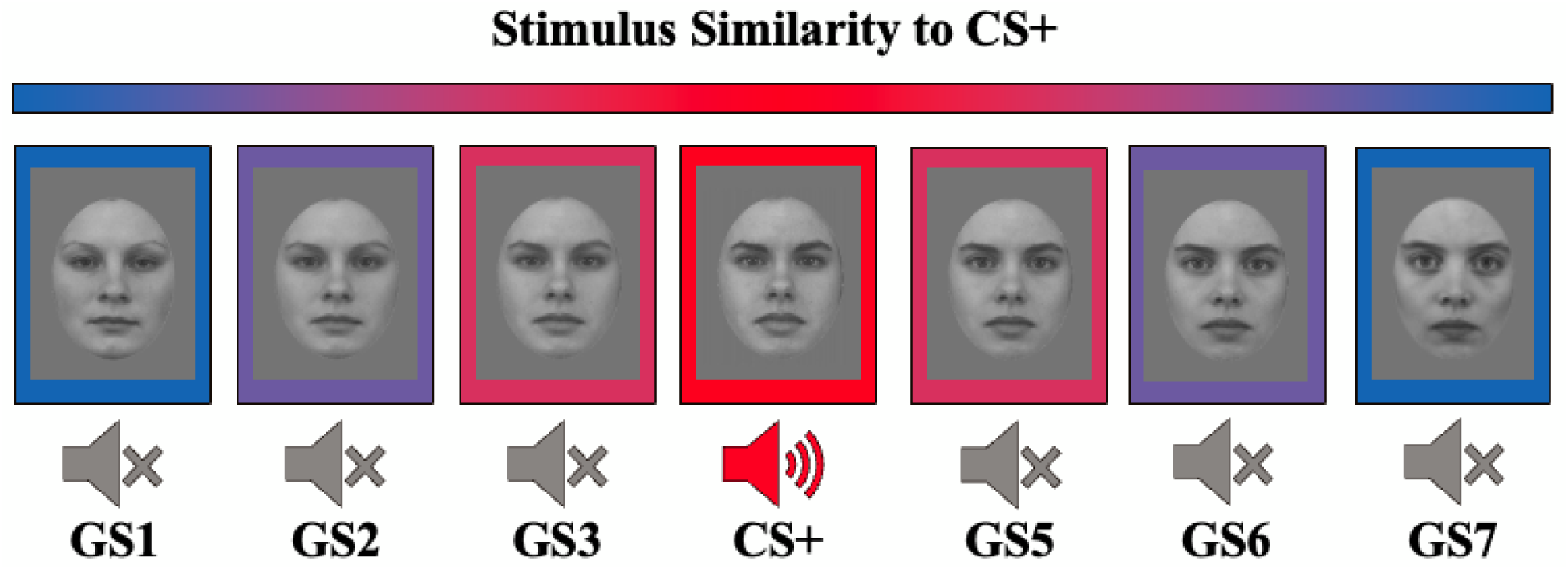
Gradient of facial stimuli. Four faces were generated using MorphX software to create systematically differing facial morphs, used as gradient stimuli (GS). The GS1, the GS7, and the conditioned stimulus (CS+) were used as the basis of the facial morphs. The GS2 and GS3 are, respectively, 30:70 and 70:30 blends of GS1 and CS+. The GS5 and GS6 are, respectively, 30:70 and 70:30 blends of CS+ and GS7. The CS+ in the middle was paired with presentation of 92 dB(A) white noise.

### Design and Procedure

During the conditioning paradigm, participants were seated in a dimly lit room (∼6 cd/m2) approximately 60cm from the eye tracker lens and approximately 120 cm from the display monitor. The paradigm (figure 3) consisted of 291 acquisition trials, in each of which one of the facial stimuli was presented in the center of the screen at visual angle of ∼7.63°. The CS+ was paired with the US at a 100% reinforcement rate. Initially, 11 booster trials (3 CS+, 4 GS1, 4 GS7) were presented in which only the CS+, GS1, and GS7 were shown to ensure familiarity with the un-morphed stimuli prior to generalization. All seven faces were pseudorandomly distributed across the remaining 280 trials. Presentation of the CS+ lasted for 3 seconds, co-terminating with a 1-second US during the last second. All other faces were only presented for 2 seconds. All faces were flickered at a frequency of 15 Hz. The intertrial interval (ITI) included the presentation of a central fixation dot. The ITI ranged from 2-4 seconds and was randomly sampled from a uniform distribution. Self-Assessment Manikins (SAMs) were presented at a visual angle of ∼21.47° at 3 fixed points: prior to conditioning (i.e. baseline), during initial acquisition (following trial 87), and during sustained acquisition (following trial 168) to collect valence and arousal ratings in response to each facial stimulus (Bradley & Lang, 1994).

**Figure 3.**
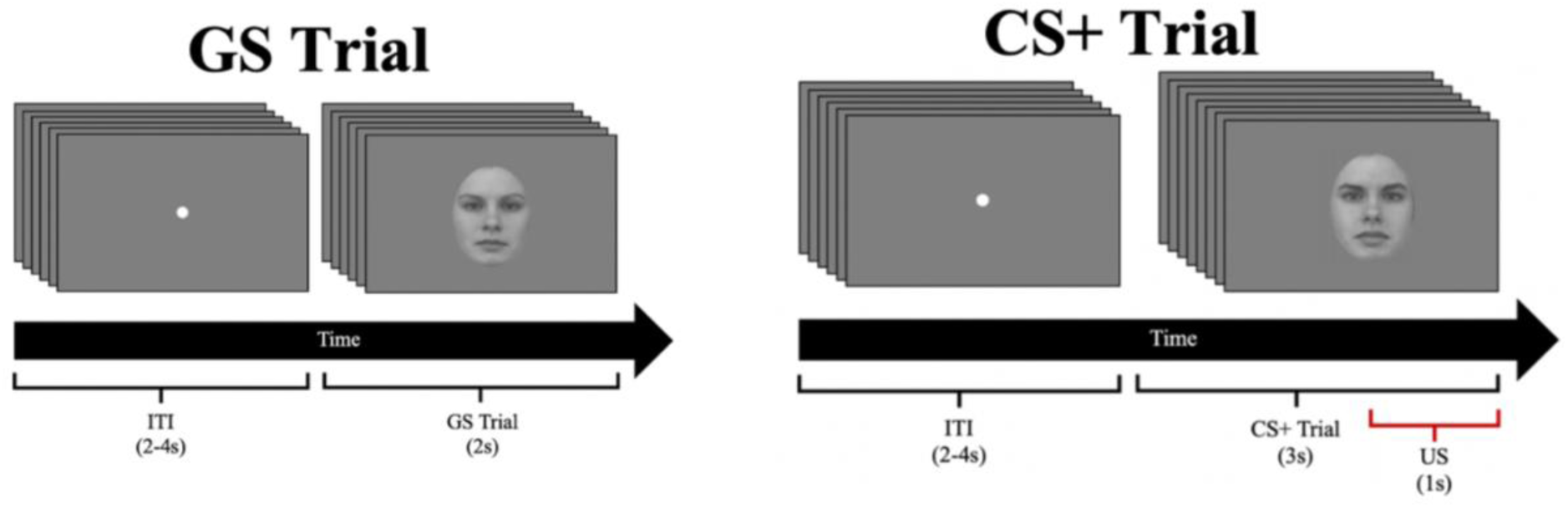
Trial structure. Stimuli were presented following a fixation dot shown during an intertrial interval (ITI) ranging from 2-4 seconds along a uniform distribution. Gradient stimuli (GS1, GS2, GS3, GS5, GS6, GS7) were presented individually in the center of the screen for 2 seconds, flickering at a driving frequency of 15Hz. The conditioned stimulus (CS+) was presented in the center of the screen for 3 seconds, co-terminating with presentation of the unconditioned stimulus (US) during the final second.

### Data Collection and Processing

A 128 channel Electrical Geodesics (EGI) electroencephalography (EEG) system (EGI, Eugene, OR) was used to record neuroelectrical signals with an input impedance of 200 MΩ. The sampling rate was 500 Hz. Impedances were kept below 60 kΩ prior to recording as recommended for EGI high-impedance amplifiers. EEG signals were referenced online to electrode Cz. EEG data were processed using Electro/MagnetoEncephalography Software (EMEGS), an open-source software implemented in MATLAB (Peyk et al., 2011). Relative to stimulus onset, data segments were extracted such that epochs were 800 ms pre-stimulus and 2200 ms post-stimulus. Data were filtered using a 40 Hz (3dB point) low pass filter (45dB/octave, 23rd order Butterworth) and a 4 Hz (3dB point) high pass filter (18dB/octave, 2nd order Butterworth). Data were re-referenced offline to the average signal across all electrodes to detect and interpolate global artifacts. Artifact rejection algorithms (per Junghöfer et al., 2000) were applied to calculate data quality indices (absolute value, standard deviation, maximum of differences across time points) for each channel and trial. A regression-based algorithm (Schlögl et al., 2007) was used to subtract activity arising from eye movements as detected using horizontal and vertical EOG channels. EEG data were then CSD-transformed using an algorithm by Junghöfer et al. (1997) to reduce the effects of volume conduction and enhance spatial specificity of the topography. The CSD/cortical mapping algorithm uses the 129-channel voltage map to calculate the current density at the level of the surface of the brain but does not provide depth information. Eye responses were recorded with an Eyelink 1000 Plus SR Research system using a 16mm lens. Segments with missing pupil data were identified and replaced via cubic spline interpolation. Only pupil data were retained and gaze data were not analyzed. For each subject, all data were separated into early and late trials using a median split and were analyzed separately.

### Self-report data

Each participant reported hedonic valence and emotional arousal in response to each face at three separate points during the paradigm. The responses from each point in the experiment across all faces were fitted via bootstrapped Bayesian evidence tests to the generalization and all-or-nothing models (see *Statistical Analysis* section).

### Pupil Analyses

Pupil dilation responses for each subject were separated into early and late trials for each participant and subtracted from an averaged baseline segment extending from –360ms to –100ms (figure 4A). The pupil data were then averaged by facial stimulus and subjected to Bayesian tests of model fit with respect to the generalization and all-or-nothing weight vectors.

**Figure 4.**
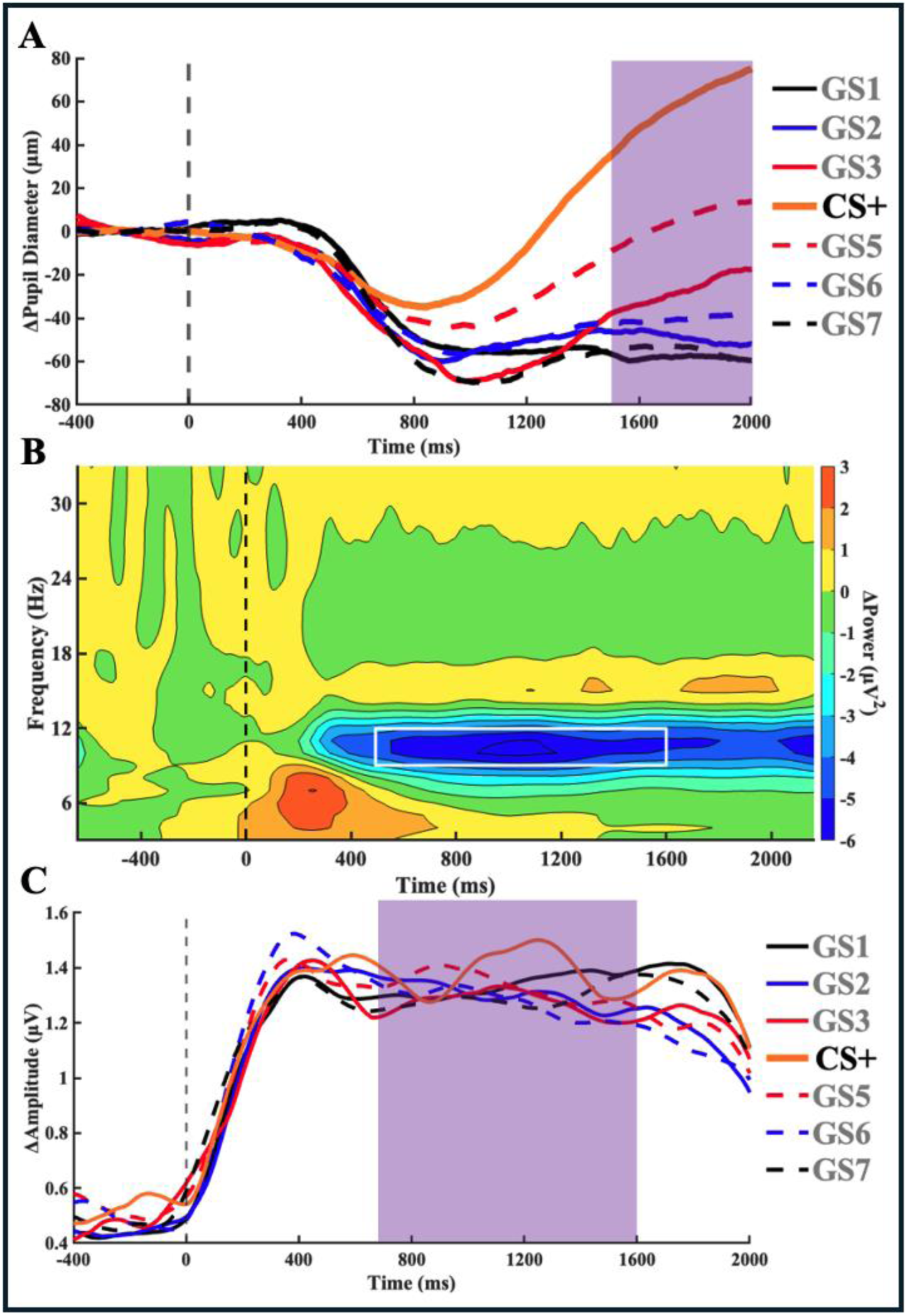
Dependent variables derived from physiological measures. A) Baseline corrected pupil size time course, averaged by condition across all participants. The purple window indicates the averaged epoch used for statistical analysis (1500ms to 2000ms). B) Baseline corrected time-frequency spectrogram from channel Oz averaged across all participants and conditions, after convolution with a family of Morlet wavelets. The white window indicates where data were averaged across frequencies (10Hz to 12Hz) and time (500ms to 1600ms) prior to statistical analysis. C) Baseline corrected time-varying 15Hz amplitude from channel Oz averaged across all participants and conditions, alongside a time-averaged topography. The purple window indicates the averaged epoch used for statistical analysis (700ms to 1600ms).

### Alpha Power Reduction

Morlet wavelet analysis was applied to the artifact-free EEG trials to measure time-varying changes in alpha (8-13 Hz) activity (figure 4B). Wavelets with central frequencies of 9.9933 Hz, 10.9927 Hz, and 11.9920 Hz was used to measure alpha-band activity. A Morlet constant of m = 7 was used as the compromise between the temporal and frequency resolutions of the wavelets. As a result of these parameters, σ_t_ = 0.1115 and σ_f_ = 1.4276 for the lowest frequency and σ_t_ = 0.0929 and σ_f_ = 1.7131 for the highest frequency. The resulting time-frequency matrices were averaged across trials and baseline-adjusted by subtracting, for each frequency, the mean power in a baseline segment from –600ms to –200ms. The convolved signals in each channel were averaged across frequencies to yield estimated alpha activity. These alpha time series were averaged across time from 500ms to 1600ms separately by facial stimulus. Because greater alpha suppression is interpreted as greater engagement with the stimulus, the weight vectors used for model fit were inverted, such that more negative values were predicted for the CS+.

### ssVEP Analyses

To isolate the time-varying 15 Hz ssVEP waveform, a Hilbert transform (9th order Butterworth) was applied to the processed EEG data after baseline subtraction using a baseline segment from –438ms to –2ms. The 15 Hz ssVEP data were separated into early and late trials, averaged across time from 700ms to 1600ms separately by facial stimulus, and subjected to model fitting procedures (figure 4C). The averaged ssVEP responses were fit to both the generalization and sharpening models. Each weight corresponds to the hypothesized ssVEP amplitude observed after presentation of each face. Model fit during these periods was determined using the bootstrapped Bayesian procedure, outlined in the subsequent section.

### Statistical Analysis

Support for the hypotheses (generalization, sharpening, all-nothing) was evaluated for all dependent variables using the a priori ideal models represented by weight vectors (figure 1). The Generalization model (M_1_) is a quadratic curve representing broad generalization. From GS1 to GS7, the values in the generalization vector were [–3.0 0.5 1.5 2.0 1.5 0.5 –3]. The all-or-nothing model (M_2_) is a function representing discrimination of the CS+ relative to the GSs. The values in the all-or-nothing model were [–1.0 –1.0 –1.0 6.0 –1.0 –1.0 –1.0]. The sharpening model (M_3_) is a difference of Gaussians, also known as a Ricker wavelet or “Mexican hat” function, which represents suppression of responses to GS’s immediately adjacent in similarity to the CS+ and an increase in physiological response as similarity to the CS+ further decreases. Values in the sharpening model were [0.5 –1.0 –2.0 5.0 –2.0 –1.0 0.5]. The inclusion of an all-or-nothing model allows for a measure of response discrimination which can be compared to the degree to which generalization occurs. Unlike the sharpening model, the all-or-nothing model is not orthogonal to generalization and as such is not a mutually exclusive hypothesis but rather an additional metric by which one can characterize these conditioned response patterns. This permits a more detailed description of observed patterns and how they change over time.

A non-parametric Bayesian approach was utilized to estimate support for fit to the ideal models. To this end, model convolution was calculated by taking the inner product of each weight vector and the sample averaged physiological or behavioral responses for each condition. As suggested by Schwarzkopf (2015), evidence supporting each model was quantified by bootstrapping inner product distributions (5000 iterations) for each model and a null distribution (calculated through permutation testing, 5000 iterations). Bayes factors (BF_M0_) were computed as the posterior odds that the inner products comprising the model distribution were greater than those of the null distribution. Greater evidence to support the null hypothesis would suggest that there is no difference in responses to any of the faces. Transitive Bayes factors were obtained to directly compare the alternative models and were computed as the posterior odds that the inner product comprising one model distribution were greater than those of the other model distribution. This bootstrapped Bayesian approach is advantageous compared to typical frequentist analyses because it is bereft of parametric assumptions about the underlying distributions, rejects the practice of delineating binary significance, and is not as susceptible to Type I error due to multiple comparisons (see Dienes, 2016). Bayes factors were log_10_-transformed to facilitate topographical mapping and to obtain a convenient and interpretable distribution. Model fits were computed separately for the initial learning phase and subsequent sustained learning phase.

## Results

### Self-reported valence and arousal

Arousal ratings (figure 5) and valence ratings (figure 6) collected at three separate occasions during the experimental session were fit to the generalization and all-or-nothing models. For baseline arousal ratings, Bayes factors indicated decisive support for the null hypothesis compared to generalization (log_10_BF_10_ = –2.13) and little evidence preferring either the null hypothesis or all-or-nothing (log_10_BF_20_ = 0.27) (table 1). Arousal ratings completed during early trials fit decisively to both of these models compared to the null (log_10_BF_10_ = 7.92, log_10_BF_20_ = 6.92). A direct comparison between generalization and all-or-nothing found insufficient evidence to prefer one over the other (log_10_BF_21_ = –0.01). Arousal ratings completed during late trials showed even greater fit to both models (log_10_BF_10_ = 12.51, log_10_BF_20_ = 16.18), and a direct comparison between alternative models showed strong evidence favoring the all-or-nothing model (log_10_BF_21_ = 1.18).

**Figure 5.**
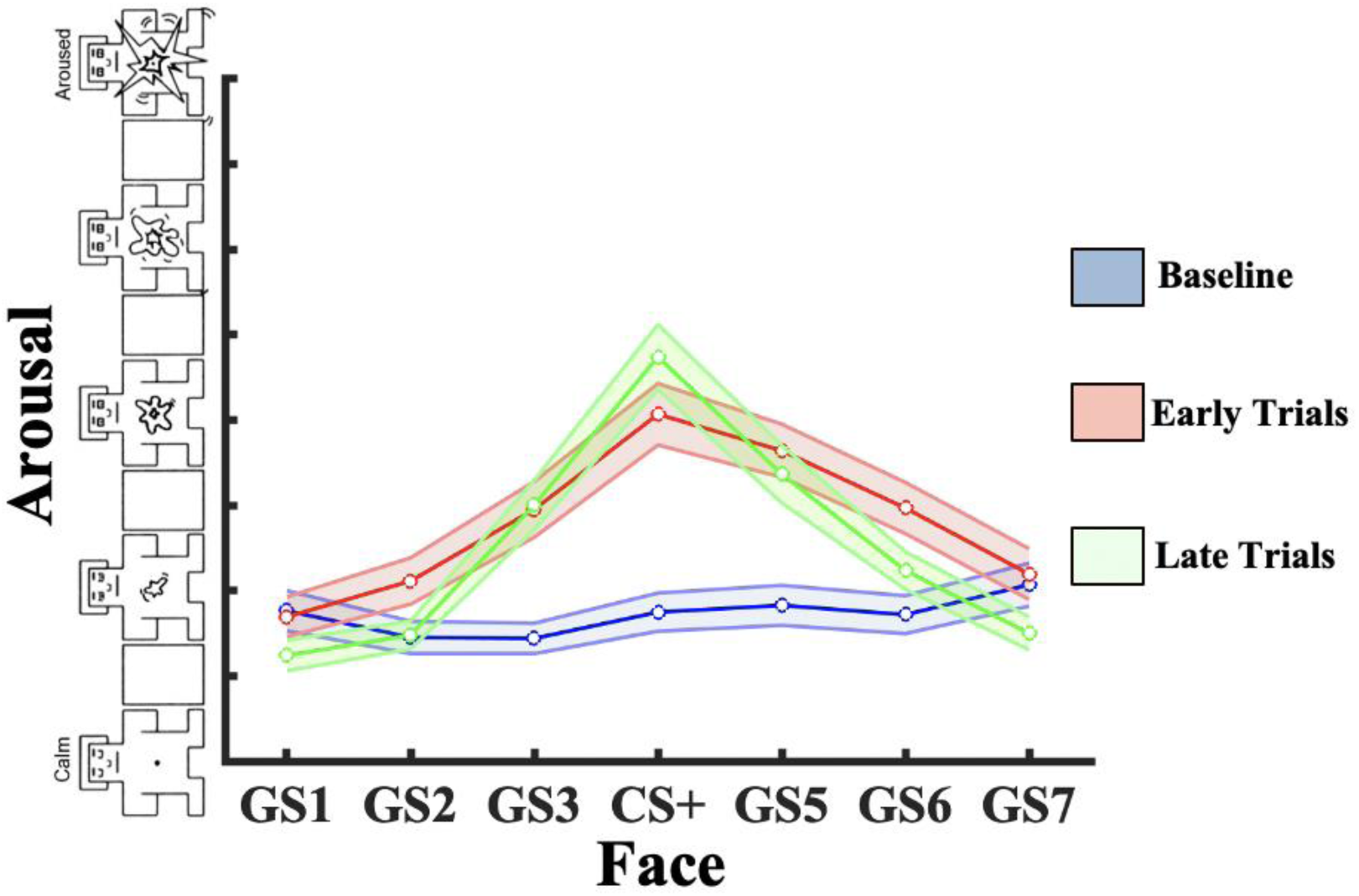
Arousal Condition Means. Arousal ratings were taken during baseline before conditioning, during the first half of trials, and during the second half of trials. An increase along the y-axis indicates an increase in reported physiological arousal or emotional intensity. SEM bars included to indicate significant differences.

**Figure 6.**
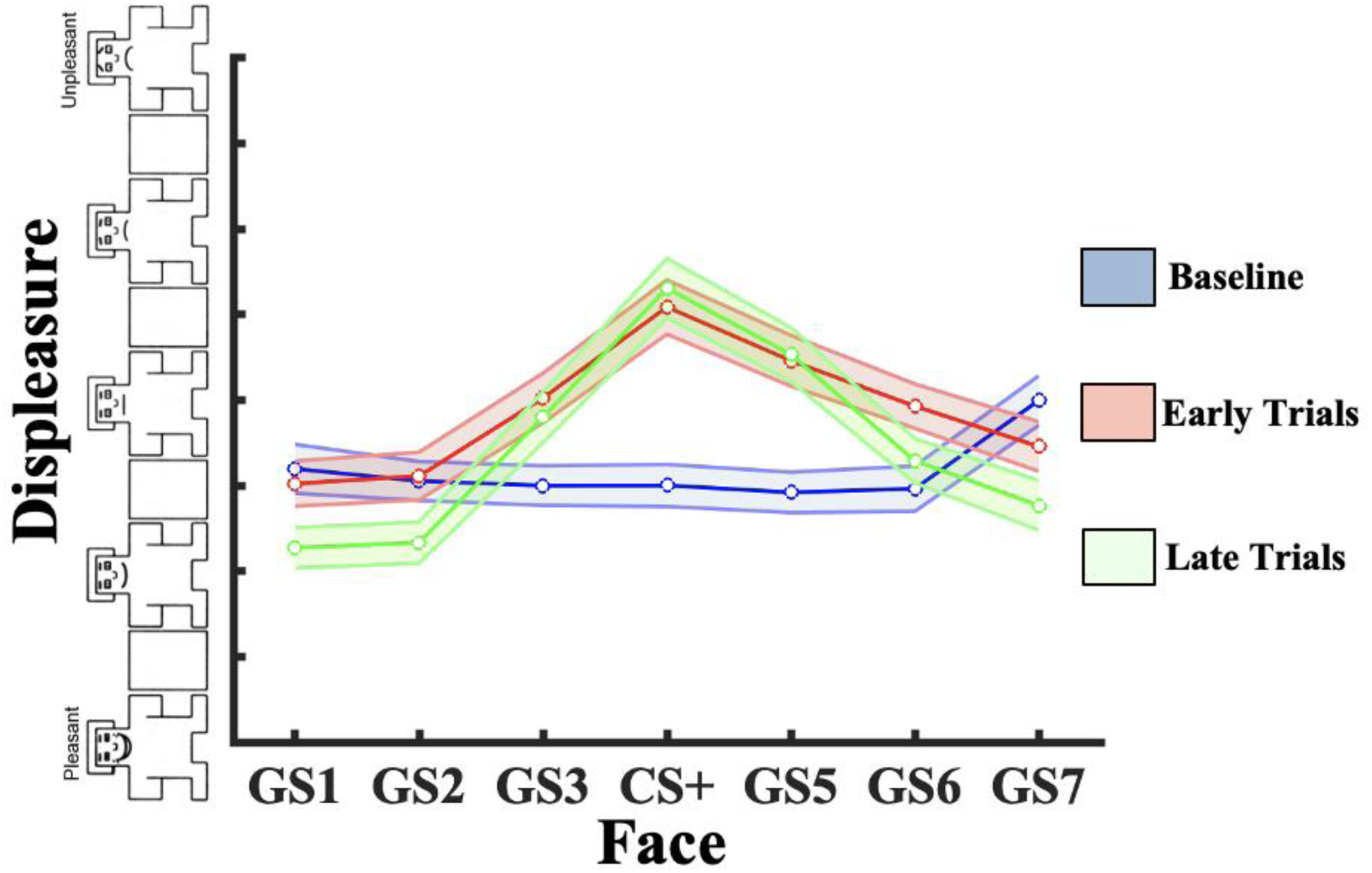
Valence Condition Means. Valence ratings were taken during baseline before conditioning, during the first half of trials, and during the second half of trials. An increase along the y-axis indicates an increase in reported negative valence or displeasure. SEM bars included to indicate significant differences.

**Table 1.** Arousal Model Fit Metrics. log_10_Bayes Factors were used to determine fit of arousal rating data to each of the hypothesized models at each rating. Direct model comparison during baseline was irrelevant to study aims and was not included.

| Model | Baseline | Early | Late |
| --- | --- | --- | --- |
| Generalization ($\log_{10}BF_{10}$) | -2.13 | 7.92 | 12.51 |
| All-or-Nothing ($\log_{10}BF_{20}$) | 0.27 | 6.92 | 16.18 |
| AON / Gen ($\log_{10}BF_{21}$) | — | -0.01 | 1.18 |

Valence ratings from baseline decisively favored the null model compared to generalization (log_10_BF_10_ = –4.36) and strongly favored the null over the all-or-nothing (log_10_BF_20_ = –1.08) (table 1). Valence ratings completed during early trials showed decisive evidence for both models (log_10_BF_10_ = 5.99, log_10_BF_20_ = 7.99), while a direct comparison between the alternative models indicated substantive evidence favoring all-or-nothing (log_10_BF_21_ = 0.69). Later valence ratings again showed an increase in fit for both models (log_10_BF_10_ = 10.36, log_10_BF_20_ = 13.02), while a direct comparison between the generalization and all-or-nothing models showed substantive support for all-or-nothing compared to generalization (log_10_BF_21_ = 0.86).

**Table 2.** Valence Model Fit Metrics. log_10_Bayes Factors were used to determine fit of hedonic valence rating data to each of the hypothesized models at each rating.

| Model | Baseline | Early | Late |
| --- | --- | --- | --- |
| Generalization ($\log_{10}\text{BF}_{10}$) | -4.36 | 5.99 | 10.36 |
| All-or-Nothing ($\log_{10}\text{BF}_{20}$) | -1.08 | 7.99 | 13.02 |
| AON / Gen ($\log_{10}\text{BF}_{21}$) | — | 0.69 | 0.86 |

### Pupil Dilation

Early average pupil responses (figure 7) decisively fit both models (log_10_BF_10_ = 4.92, log_10_BF_20_ = 7.77), while a direct comparison showed no preference for either (log_10_BF_21_ = –0.03) (table 3). During late trials, average pupil dilation patterns showed greater fit to both models (log_10_BF_10_ = 5.24, log_10_BF_20_ = 9.70), while a direct comparison between them found strong support for the all-or-nothing model over the generalization model (log_10_BF_21_ = 1.61).

**Figure 7.**
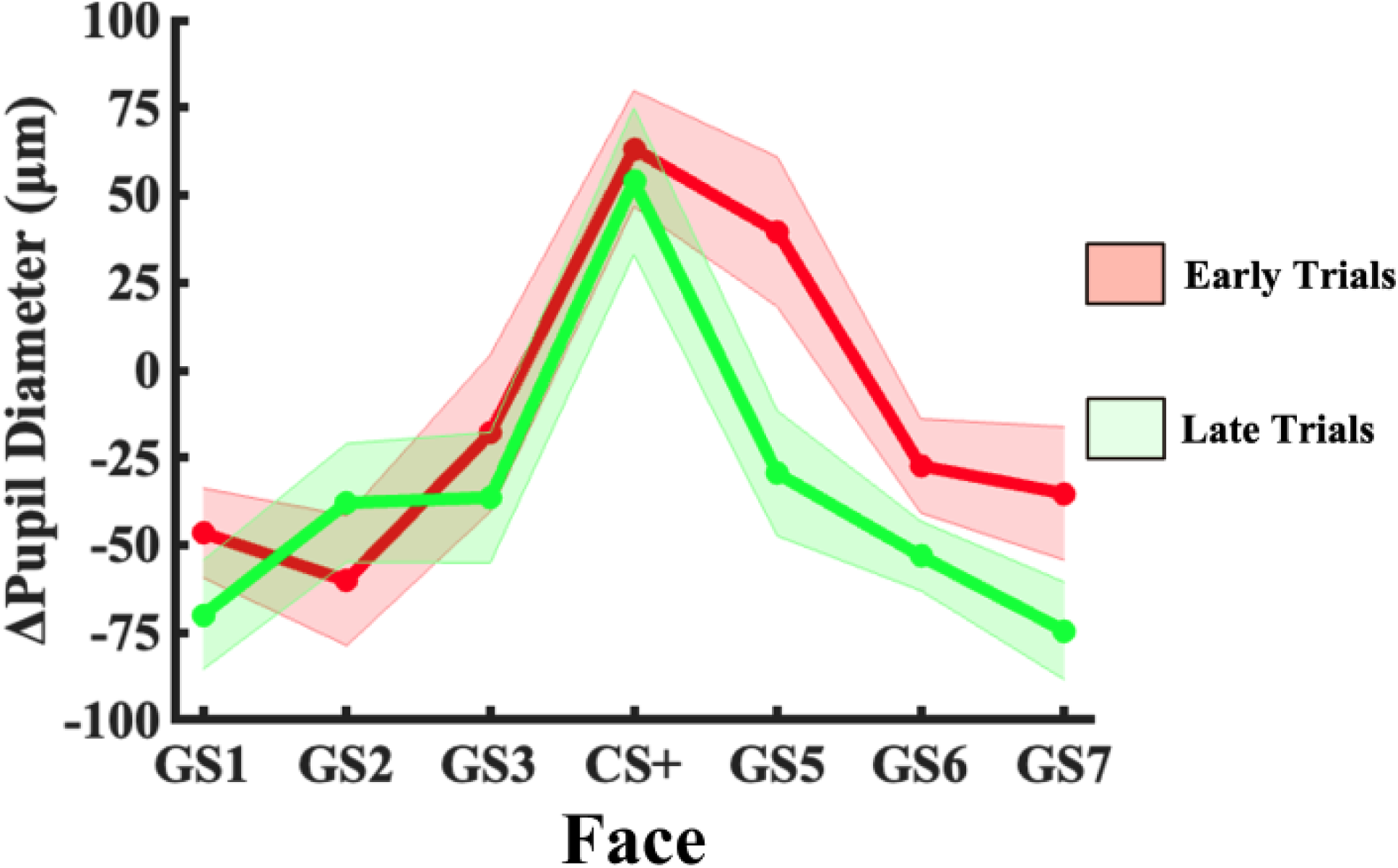
Averaged Pupillary Response Condition Means. Pupil responses in each condition averaged across the analytic epoch (1500ms-2000ms) and across subjects. Shaded error bars depicting the within-subjects standard error of the mean (SEM) included to indicate condition differences.

**Table 3.** Pupillary Responses Model Fit Metrics. log_10_Bayes Factors were used to determine fit of pupillary responses across conditions to each of the hypothesized models at each rating.

| Model | Early | Late |
| --- | --- | --- |
| Generalization ($\log_{10}\text{BF}_{10}$) | 4.92 | 5.24 |
| All-or-Nothing ($\log_{10}\text{BF}_{20}$) | 7.77 | 9.70 |
| AON / Gen ($\log_{10}\text{BF}_{21}$) | -0.03 | 1.61 |

### Alpha Suppression Analyses

Patterns of alpha suppression showed strong to decisive fit to the all-or-nothing model compared to the null in bilateral frontal (F4: log_10_BF_10_ = 0.81, log_10_BF_20_ = 3.41, log_10_BF_12_ = – 1.53) and medial centroparietal channels (CPz: log_10_BF_10_ = 0.21, log_10_BF_20_ = 2.11, log_10_BF_12_ = – 1.35) during early trials (figure 8). There was also decisive fit to generalization in right centroparietal channels (CP6: log_10_BF_10_ = 2.50, log_10_BF_20_ = .23, log_10_BF_12_ = 1.50). During late trials, there was widespread decisive fit to both models in occipitotemporal channels, but the data best fit the all-or-nothing model in right occipitotemporal channels (P10: log_10_BF_10_ = 2.60, log_10_BF_20_ = 4.57, log_10_BF_12_ = –0.87). Additionally, medial centroparietal channels decisively fit the all-or-nothing model while showing strong support for fit to the generalization model (channel 54, near CPz: log_10_BF_10_ = 1.50, log_10_BF_20_ = 3.65, log_10_BF_12_ = –1.27). Figure 9 shows the average condition means of notable channel clusters based on model fit results.

**Figure 8.**
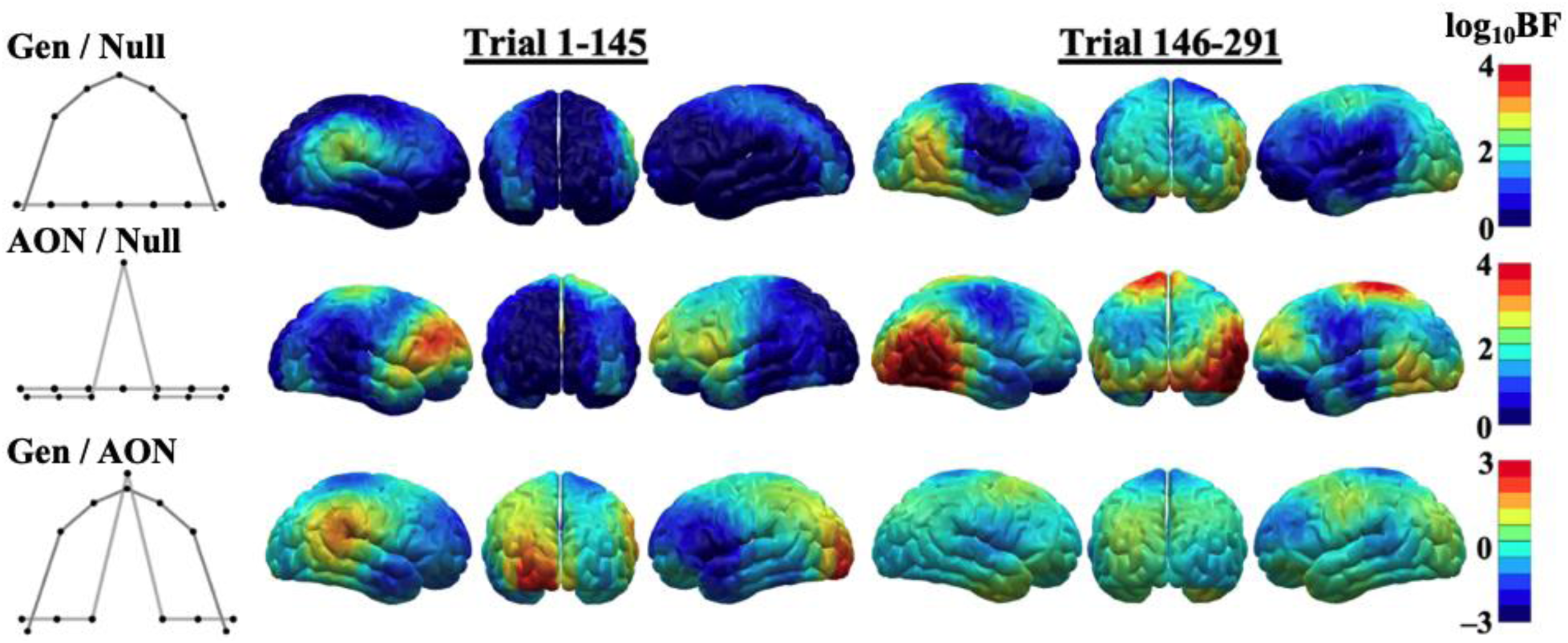
Brain Mapping of Alpha Suppression Bayesian Model Fits. log_10_Bayes Factors were computed comparing fit of electrocortical responses to each hypothesized model to the null, alongside a direct comparison of model evidence between both models. This was done for all channels. Topographies map model fits at each chain to their corresponding cortical region. Results show widespread overlap of fit to both models, and general increase in model fit over time. Strongest model fit to both models occurs during later trials in right occipitotemporal and central regions.

**Figure 9.**
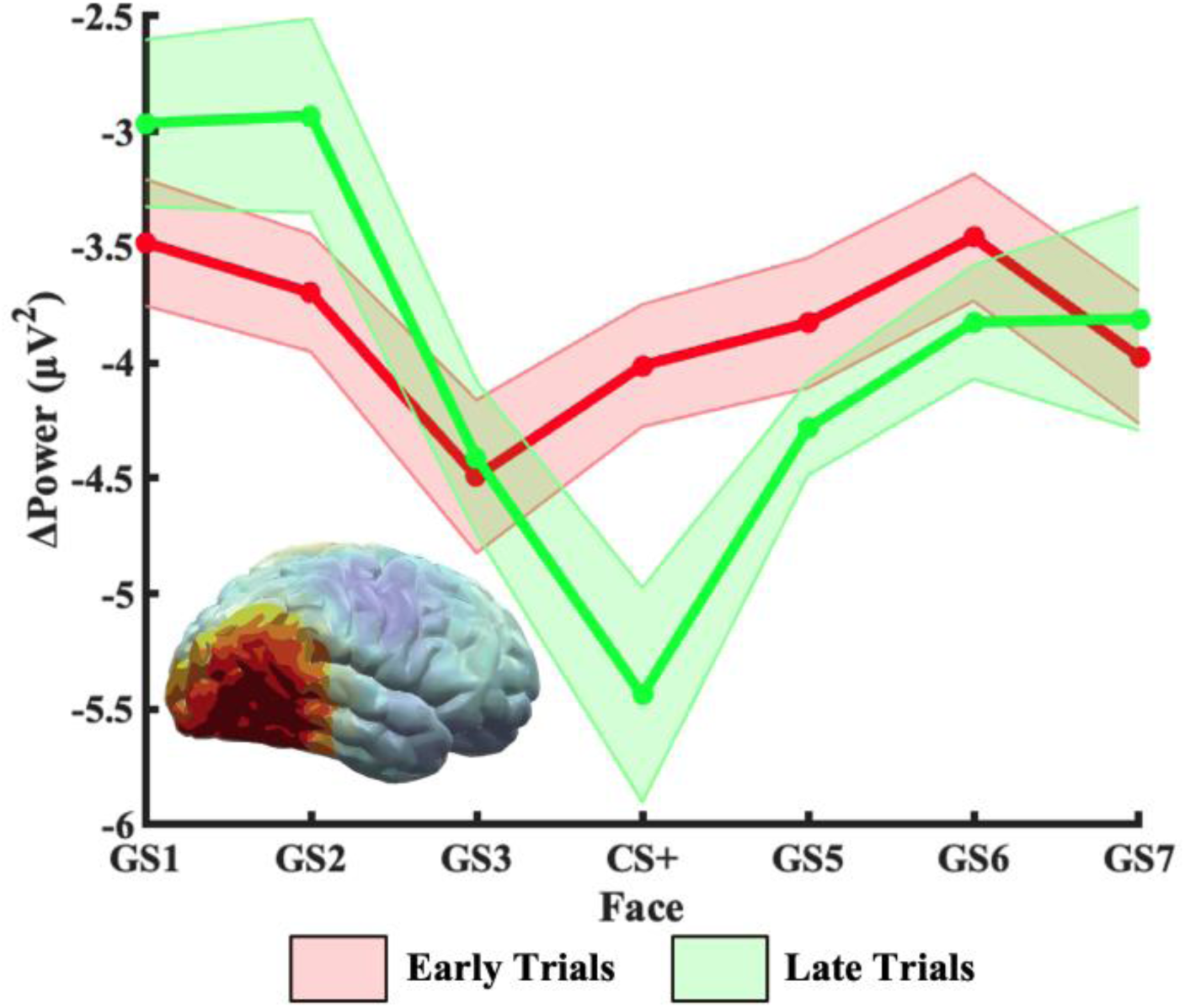
Averaged Alpha Power Condition Means. Baseline-corrected alpha power was averaged across subjects and across the analytic epoch (500ms-1600ms). Additionally, power was averaged across a 5-channel right occipitotemporal cluster showing greatest model fit to the all-or-nothing model to demonstrate changes in cortical tuning with respect to the stimulus gradient. Shaded error bars depicting the within-subjects standard error of the mean (SEM) included to indicate condition differences.

### ssVEP Analyses

Bayesian tests of model fit to average ssVEP responses indicated decisive model fit to sharpening in medial occipitoparietal (PO3: log_10_BF_10_ = –0.05, log_10_BF_30_ = 2.96, log_10_BF_13_ = – 2.36) and right prefrontal channels (AF4: log_10_BF_10_ = 1.46, log_10_BF_30_ = 4.02, log_10_BF_13_ = –1.49) during early trials (figure 10). Additionally, during early trials there was decisive fit to generalization in bilateral occipital (PO7: log_10_BF_10_ = 2.21, log_10_BF_30_ = –0.71, log_10_BF_13_ = 2.41), right centroparietal (PO7: log_10_BF_10_ = 2.21, log_10_BF_30_ = –0.71, log_10_BF_13_ = 2.41), and left frontal channels (FC1: log_10_BF_10_ = 2.35, log_10_BF_30_ = 1.23, log_10_BF_13_ = 0.52). During late trials, there was greater fit to generalization in left frontal channels (FC1: log_10_BF_10_ = 3.37, log_10_BF_30_ = 0.68, log_10_BF_13_ = 1.76).

**Figure 10.**
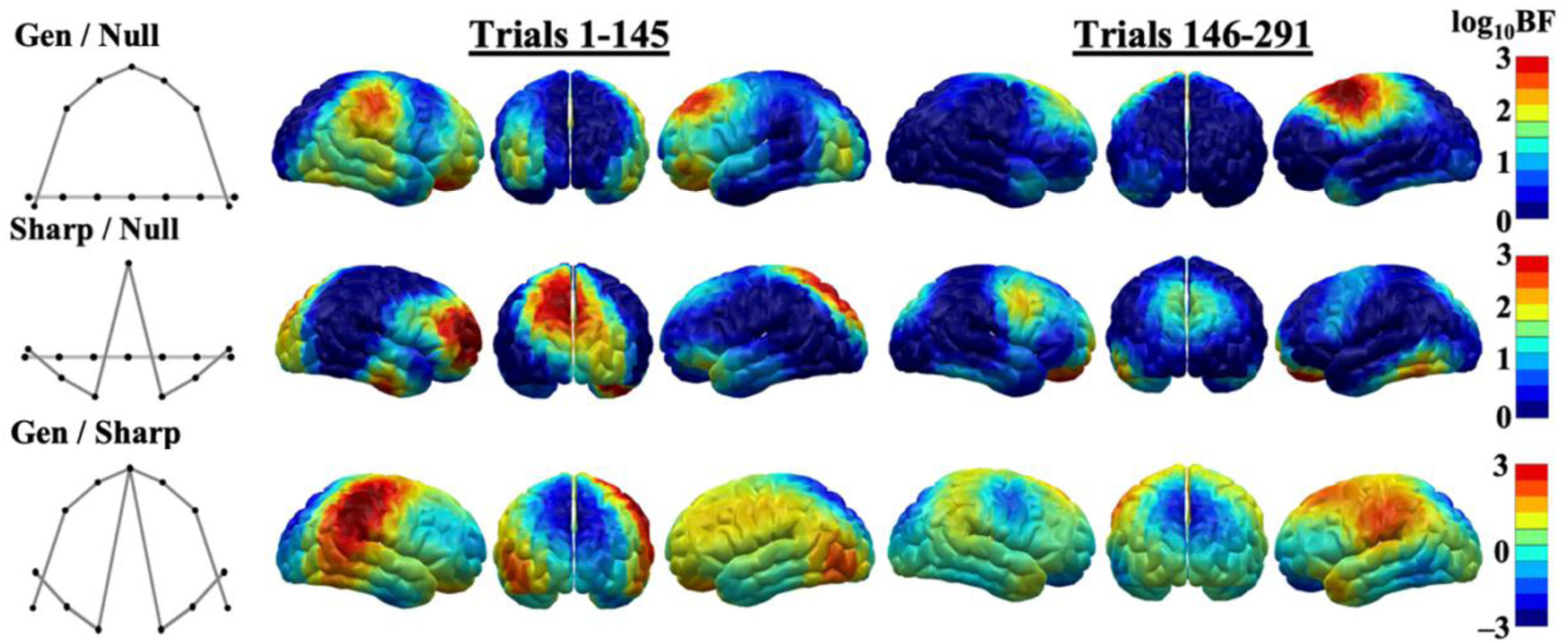
Brain Mapping of ssVEP Bayesian Model Fits. log_10_Bayes Factors were computed comparing fit of electrocortical responses to each hypothesized model to the null, alongside a direct comparison of model evidence between both models. This was done for all channels. Topographies map model fits at each channel to their corresponding cortical region. Results show large differences in model fit between conditioning periods. Most notable is the occipitoparietal channels, where fit to the sharpening pattern is decisive during early trials but decreases to strong in later trials.

**Figure 11.**
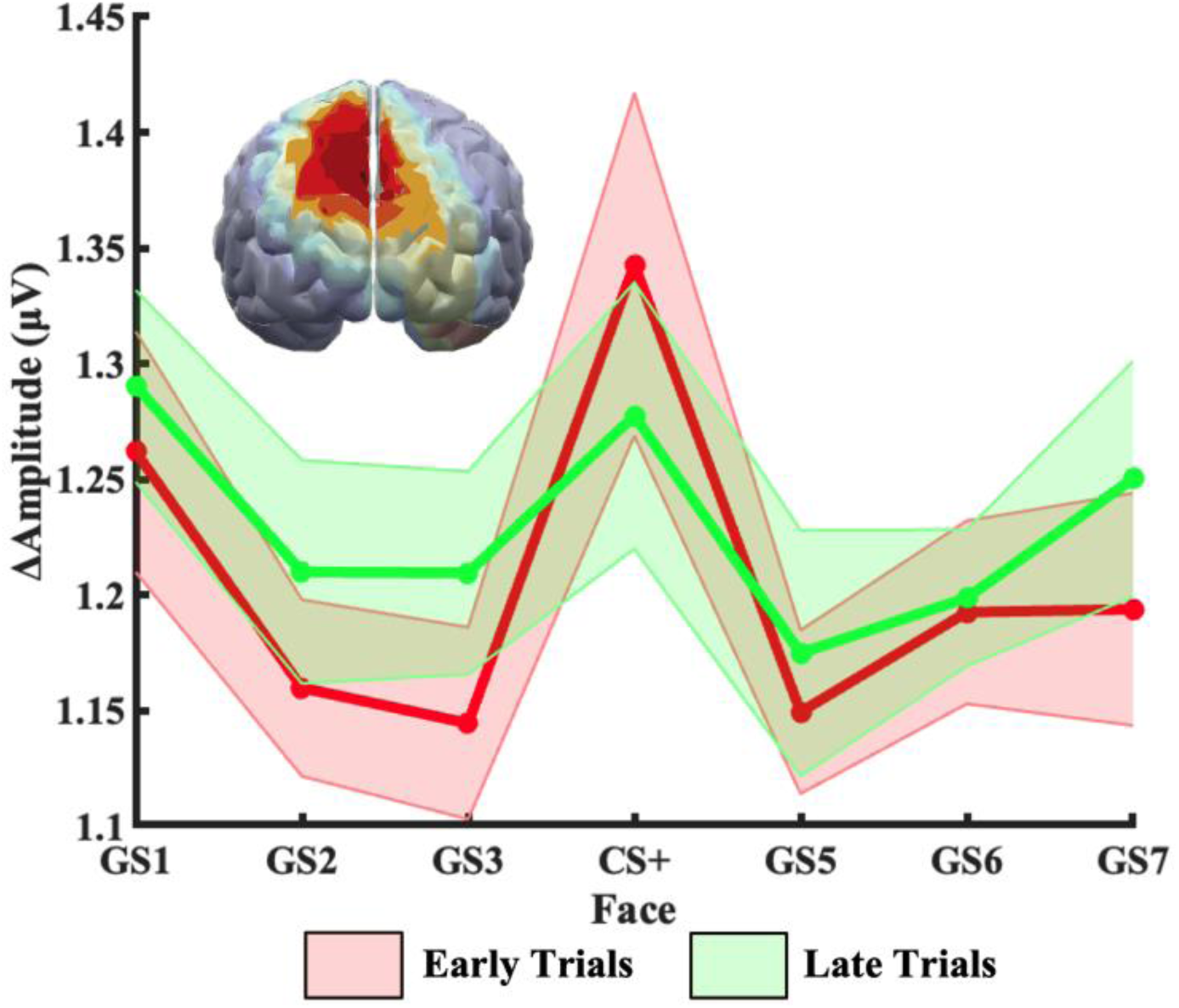
Averaged Steady-State Visual Evoked Potential (ssVEP) Condition Means. Baseline-corrected ssVEP amplitude was averaged across subjects and across the analytic epoch (700ms-1600ms). Additionally, amplitude was averaged across a 5-channel occipitoparietal cluster showing greatest model fit to the sharpening model to demonstrate changes in cortical tuning with respect to the stimulus gradient. Shaded error bars depicting the within-subjects standard error of the mean (SEM) included to indicate condition differences.

## Discussion

Generalization learning shapes the tuning functions of response variables indexing a range of physiological processes. Here we asked to what extent known generalization tuning profiles would undergo further changes as learning progresses beyond a typical trial count of 10 to 20 trials per condition. Stegmann and colleagues (2020) found a sharpening pattern in visuocortical areas when presenting 15 generalization trials per condition, after an initial conditioning regimen with 15 trials for CS+ and CS-only. Their finding, along with several others (see Li & Keil, 2023 for a review), is consistent with the idea that neurons in visual cortex that are tuned to similar features as a CS+ will be subject to heightened suppression, presumably through lateral inhibitory interactions with the strengthened CS+ representation. Using 7 levels of generalization, the present study also adds refined tuning pattern information, compared to other reports (e.g., Friedl & Keil, 2021). The present study replicated the finding by Stegmann and colleagues (2020), but also provides important information regarding the changes during prolonged, sustained learning. Specifically, after 40 generalization trials per condition, visuocortical areas still displayed sharpening, but to a substantially lesser extent than during initial learning. Other variables also showed evidence of continued retuning as learning progressed: For example, although generalization tuning of pupillary responses was present in early and late trials, these response patterns showed an increased tendency toward discrimination (all-or-nothing tuning) rather than greater generalization during the latter part of acquisition. A similar trend was observed for arousal ratings, and for alpha power suppression, both of which showed increased all-or-nothing tuning. We discuss the implications of these findings in the following, organized by dependent variable.

### Self-reported valence and arousal

In line with a body of prior studies examining conditioned fear generalization (Stegmann et al., 2020; Ahrens et al., 2016), the participants in the present study were able to correctly identify the CS+ face amid an array of similar faces. As stated above, the gradient of negative valence and arousal to GS’s as a function of CS+ similarity became steeper over time, increasingly resembling an all-or-nothing tuning function. Post-hoc t-tests comparing early and late trials highlight this observation. Faces with less similarity to the CS+ (GS1, GS2, GS6, GS7) decreased in negative valence and emotional arousal (p < 0.05), while proximal faces (GS3, GS5) did not (p > 0.05). Despite this trend, the data were still described by a generalization function, albeit a narrower one than prescribed by our a priori models. Ongoing work from our laboratory aims to develop methods for more precisely and continuously characterizing tuning profiles, avoiding a priori models altogether.

Faces used in this study were developed in pilot studies, to ensure discriminability and neutral affect. Despite these efforts, it is apparent that participants in the present study rated GS7 as significantly more unpleasant than the other faces at baseline. This is consistent with verbal reports during debriefing from a sizeable subset of participants who commented that GS7 was least attractive and least pleasant. Unlike simple stimuli such as shapes and Gabor patches, faces may trigger unpleasant or pleasant emotional reactions prior to conditioning. Faces with neutral expressions elicit social and emotional trait attributions (Said et al., 2009). This presents a possible confounding effect on self-report responses and could also affect other measures of conditioned fear. Future studies may collect ratings in terms of valence and arousal for various facial gradients presented to participants without any conditioning to reduce this confound.

Relatedly, a shortcoming of the present study with regard to external validity is the lack of diversity in both the chosen stimuli and participants recruited. Approximately 83% of the participants who completed the study were white, 82% were female, and 73% were both white and female. Abundant work in social psychology and social neuroscience has shown that face perception is affected by both in-group and out-group biases based on sex and ethnicity. These biases have been shown to influence participants’ appraisal of trustworthiness, happiness, and threat associated with a given face (Schmid et al., 2022; Kawakami et al., 2014; Friesen et al., 2019; Penton-Voak et al., 2001). Particularly germane to this study is the classic finding of an “own-race bias” where participants are better able to encode and discriminate faces that appear to share their race (Meissner & Brigham, 2001; Goldinger et al., 2009). Future work ought therefore not only recruit a more diverse sample but also utilize a more diverse stimulus set.

### Pupil Dilation

Pupil response tuning changed substantially during late trials relative to early trials. Early pupil responses displayed, as expected, a broad generalization pattern as reported previously (e.g., Friedl & Keil, 2020). However, while generalization still occurred during late trials, there was a shift toward discrimination of the CS+ and heightened all-or-nothing tuning. Comparable studies (Roesmann et al., 2022; Reutter & Gamer, 2023) have also reported generalization of pupil dilation across the conditioned stimulus gradient, but did not report on the temporal evolution and extent of this tuning pattern as learning progresses. These results suggest that sympathetic responses reflect response generalization in which output systems display defensive dispositions that are scaled relative to their similarity to the conditioned threat cue. In evolutionary terms, this is adaptive because showing potentially false-positive responses to potential threat often possess favorable cost versus benefit ratios, in the context of threat and danger. Specifically, false-positive defensive responses are less costly than false-negative responses (failing to respond), when the organism is threatened. For the same reasons, generalization tends to be stronger in situations characterized by uncertainty or by diffuse threat. In the present study, this was reflected in initially more pronounced generalization, followed by narrower tuning later in the experimental session, when participants were more certain about the contingencies. Of note, the similarity between self-reported emotional arousal and pupil dilation is consistent with similar findings, arguing that both involve engagement of the sympathetic nervous system (Bradley et al., 2008).

### Alpha Power Reduction

Alpha suppression was greatest primarily in parietal and occipitotemporal areas and showed a strong preference for the right hemisphere. Alpha suppression was also prominent in centroparietal channels, near the vertex. Over the course of conditioning, alpha suppression responses showed a generalization pattern that was most pronounced in right occipitotemporal and frontal regions. Right lateralized brain responses to faces are consistent with work on face perception, and are thought to indicate right lateralized face-specific areas, including the Fusiform Face Area (FFA) and the Occipital Face Area (OFA), which are more prominent in the right hemisphere (Rossion et al., 2003; Rossion, 2014; Fisher et al., 2016).

Paralleling pupil responses, alpha power changes showed a notable shift towards all-or-nothing tuning in late trials. This is consistent with earlier work with spatial conditioning (Friedl & Keil, 2020), which also found generalization during early trials. It is also consistent with alpha power changes with generalization conditioning in the auditory domain (Farkas et al., 2023). When compared to ssVEPs, discussed below, Bayes factors overall were substantially greater when applied to alpha power changes, highlighting the robust nature of alpha power as a metric of aversive conditioning (Bacigalupo & Luck, 2022).

### ssVEP

Occipitoparietal steady-state responses to the similarity gradient resembled the familiar sharpening, or “Mexican hat,” pattern in visual areas found in previous studies (McTeague et al., 2015; Stegmann et al., 2020). This sharpening pattern became attenuated during the latter half of trials. In other words, during early trials, while visuocortical neurons did maximally respond to the CS+, responses to stimuli adjacent in similarity (GS3 and GS5) were suppressed. Localization of this pattern differed from some prior studies in that it was more anterior, showing a maximum in occipitoparietal regions rather than lower primary visual areas. As noted in other research (Li & Keil, 2023) early sharpening is in strong contrast with the broad generalization and all-or-nothing patterns exhibited by self-report, pupillometry, and alpha data. Work by Petro et al. (2017) supports the hypothesis that the increased responsiveness of the ssVEP to CS+ relative to CS– cues results from re-entrant signals originating in anterior regions, which modulate the sensitivity of visuocortical neurons—a notion that was also supported by inter-site phase locking analyses in McTeague et al. (2015). It is possible that the sharpening pattern observed in the present study is produced by a similar re-entrant mechanism, but future work utilizing simultaneous EEG-fMRI is necessary to corroborate this notion.

Unlike alpha suppression, ssVEP signals are thought to index engagement of lower-tier visuocortical processes (Wieser et al., 2016; Bazanova & Vernon, 2014). Neurophysiological work in cats suggests that, during perception of a given simple orientation, horizontal corticocortical inputs tuned to similar orientations inhibit input from the lateral geniculate nucleus (LGN; Sengpiel et al., 1997; Gilbert & Wiesel, 1989). This increases the selectivity of orientation-specific neurons being directly excited by an orientation and exaggerates the differences between similar orientations. On a broader systems level, Hochstein & Ahissar (2002) proposed that after initial recognition of a visual stimulus, top-down signals are propagated toward lower visual areas to enable processing of finer visual details. Similar notions may apply to the functional relevance of sharpened tuning as observed here, ultimately heightening the contrast between a threat-associated and a similar non-threatening person or face. Such contrast has been argued to be adaptive as it heightens accurate discrimination of threat and safety at an early level of perceptual processing, supporting efficient and rapid identification of relevant cues (Li et al., 2019; Pourtois et al., 2013).

Contrary to our hypotheses, decisive fit to both models was found in motor and frontal regions. This is unexpected since average topographies localize ssVEP responses to occipital and occipitoparietal regions. Additionally, multiple studies have localized the source of ssVEP signals to visual cortex (Petro et al., 2017; di Russo et al., 2007; Andersen et al., 2009). One limitation of the present approach is that fit to model weights was based on how well relative magnitude of responses fit pre-defined patterns with no absolute magnitude threshold such as amplitude or signal-to-noise ratio. With this in mind, there is no clear explanation for the observed motor and frontal ssVEP model fits and thus these results should be interpreted cautiously.

## Conclusion

The current study showed that visuocortical tuning functions prompted by aversive generalization learning continue to change as learning progresses beyond an initial learning stage. Visuocortical tuning starkly contrasts with the other measures, exhibiting a sharpening pattern rather than generalization tuning. Selective alpha reduction showed generalization and all-or-nothing tuning in face-selective cortical tissue. Together, the present data support the idea that sustained experience with an aversive cue prompts continuous processes of optimizing perception and attention, accompanied by changes in the large-scale configuration of the brain regions involved in defensive processing. Future work will utilize multi-modal analyses to examine the functional interactions between affect-biased attentional systems and defensive motor responses.

## Acknowledgments

The aforementioned research was supported by grant #N00014-21-1-2324 from the Office of Naval Research, and R01MH125615 from the National Institute of Mental Health.

